# ChromFormer: A transformer-based model for 3D genome structure prediction

**DOI:** 10.1101/2022.11.15.516571

**Authors:** Henry Valeyre, Pushpak Pati, Federico Gossi, Vignesh Ram Somnath, Adriano Martinelli, Maria Anna Rapsomaniki

## Abstract

Recent research has shown that the three-dimensional (3D) genome structure is strongly linked to cell function. Modeling the 3D genome structure can not only elucidate vital biological processes, but also reveal structural disruptions linked to disease. In the absence of experimental techniques able to determine 3D chromatin structure, this task is achieved computationally by exploiting chromatin interaction frequencies as measured by high-throughput chromosome conformation capture (Hi-C) data. However, existing methods are unsupervised, and limited by underlying assumptions. In this work, we present a novel framework for 3D chromatin structure prediction from Hi-C data. The framework consists of, a novel *synthetic data generation module* that simulates realistic structures and corresponding Hi-C matrices, and ChromFormer, a transformer-based model to predict 3D chromatin structures from standalone Hi-C data, while providing local structural-level confidence estimates. Our solution outperforms existing methods when tested on unseen synthetic data, and achieves comparable results on experimental data for a full eukaryotic genome. The code, data, and models can be accessed at https://github.com/AI4SCR/ChromFormer.

## 1 Introduction

A plethora of recent studies have illustrated that the 3D genome organization affects if, when, and how genetic information is expressed [2] to ensure controlled execution of essential cellular processes [20, 19]. As genome misfolding is increasingly linked to different diseases [5, 13], modeling the 3D structure of chromatin can not only provide mechanistic insights on vital cell processes, but also identify structural disease biomarkers [1]. However, there exists no method to experimentally determine 3D genome structure. Chromatin organization is studied implicitly using high-throughput chromosome conformation capture (Hi-C) experiments [12] that produce a *contact map* containing interaction frequencies among tiny DNA fragments (loci) across the genome. In absence of experimental methods, a plethora of computational approaches that infer 3D chromatin structure from Hi-C contact maps have emerged [17]. Most methods map interaction frequencies to 3D Euclidean distances via a parametric *transfer function* and use them as constraints to solve an optimization problem [18]. However, the transfer function differs between organisms and resolutions [26]. Recently, data-driven methods employing manifold learning to learn a 3D representation of Hi-C data without a transfer function assumption have emerged, such as GEM [27]) and REACH-3D [3]. Still, in the absence of ground truth, the learning capacity of these unsupervised methods is limited, as their architecture is not designed to generate encodings resembling 3D structures. In this work we propose a novel framework for predicting 3D chromatin structure, inspired by TECH-3D, a state-of-the-art transfer learning method [6]. In contrast to TECH-3D, we propose a synthetic data generation module that simulates biologically-informed 3D structures and corresponding Hi-C matrices using optimal transport (OT), and ChromFormer, a transformer-based model that predicts 3D structures from Hi-C matrices, while also estimating the confidence of the prediction.

## 2 Methods

Our proposed framework, presented in Figure 1, consists of a *synthetic data generation* component that simulates paired chromatin structures and Hi-C matrices, and ChromFormer, a transformerbased model that is trained on the synthetic data and learns to map Hi-C matrices to 3D chromatin structures. ChromFormer also outputs loci-level confidence values using a calibration network that corrects the predicted confidence logits to estimated confidence values.

**Figure 1:**
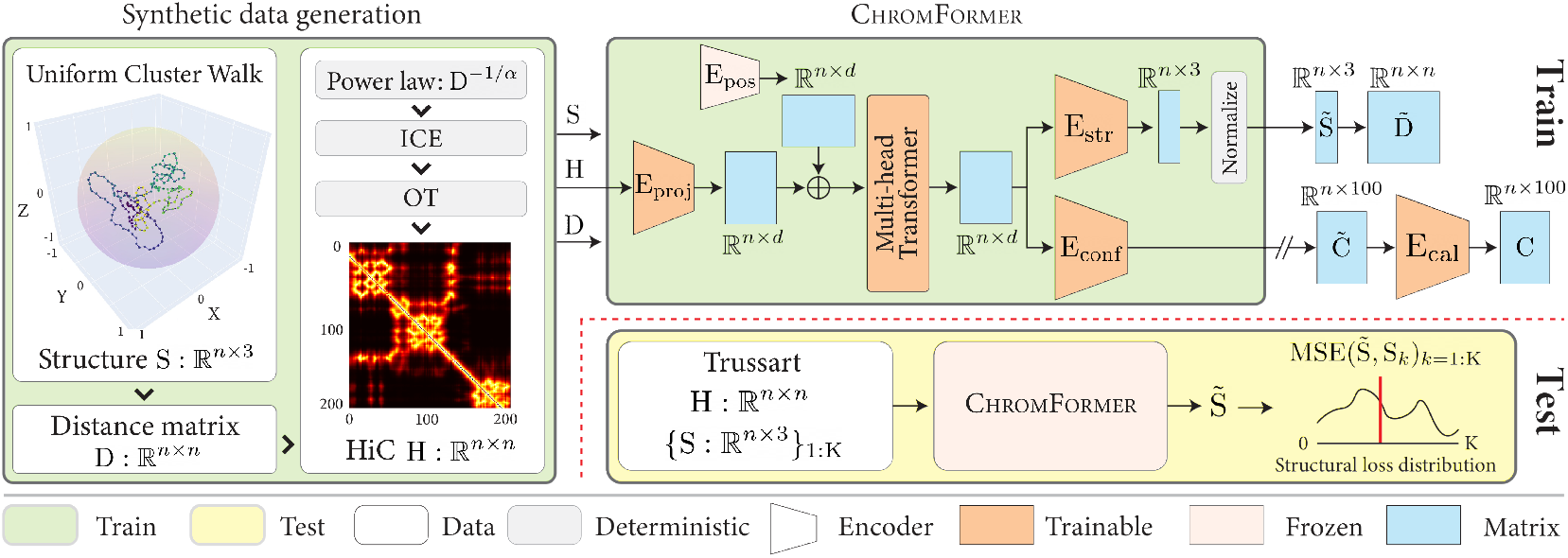
Overview of our proposed chromatin structure prediction framework.

### Synthetic Data Generation

To address the lack of ground truth data, in our framework we first propose a Uniform Cluster Walk to generate pairs of synthetic 3D chromatin structures and Hi-C matrices. To resemble real 3D chromatin structures, the synthetic structures need to satisfy the following constraints: i) smoothness and compactness, ii) equidistance of consecutive loci, and iii) presence of clustered regions resembling topologically associated domains (TADs) [16]. To this end, given a number of *n* loci, we generate a structure S ∈ ℝ^n×3^, while employing rejection sampling at different levels to ensure biological plausibility. More formally, for a given locus 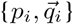, we first sample *r* ∼ 𝒰 (1, −1) where ||*r*||_2_ ≤1 to ensure isotropy. 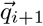 is then chosen as 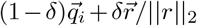, where *δ* is a smoothness parameter, followed by *L*_2_ normalization. The next locus *p*_*i*+1_ is derived as 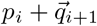. To ensure S is compact, *p*_*i*+1_ is subject to rejection sampling according to a Gaussian 𝒩 (*p*_0_, *σ*), where *p*_0_ is the starting locus and *σ* controls the degree of compactness. *p*_*i*+1_ is rejected if,

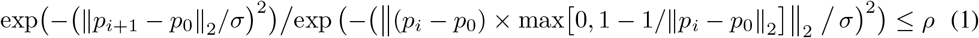

where, *ρ* is a sampled probability. Further, we create a TAD of length l << *n*, starting at *p*_*i*_, by using rejection sampling with a Gaussian 𝒩 (*p*_*i*_, *ν*) and Equation 1. As TADs are more compact, we set *ν < σ*. We enforce a distance of at least *k* loci after a TAD ending before being eligible to form another TAD. More details on the structure generation are given in the Appendix A.1. S is then centered and normalized by the maximum *L*_2_ norm of loci in S. The corresponding distance matrix D and Hi-C matrix H are generated using pairwise Euclidean distances between all loci in S, and a power law D^−1*/α*^, respectively, where *α* is a tunable parameter. We normalize H using iterative correction and eigenvector decomposition (ICE) normalization [14] and employ optimal transport (OT) [25] to match the synthetic to real Hi-C matrix. First, the real Hi-C is min-max normalized, and then OT is applied to transform the frequency distribution of the values in the synthetic Hi-C matrix to the frequency distribution of the values in the real Hi-C matrices (more details in the Appendix A.2). Notably, the usage of OT mitigates the distribution gap without any training and without requiring the real chromatin structures.

### ChromFormer

The synthetic data generator outputs sets of three matrices, S ∈ ℝ^n×3^, D ∈ ℝ^n×n^, and H ∈ ℝ^n×n^. ChromFormer operates on H to predict 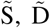 and confidence 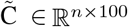. ChromFormer builds on the hypothesis that the relative spatial distribution influences the formulation of a 3D chromatin structure, and we use a Transformer [24] to exploit this hypothesis. First, ChromFormer employs *E*_proj_ : ℝ^n×n^ → ℝ^n×d^ to project H onto lower dimensional embeddings E ∈ ℝ^n×d^, *d < n*. E is summed with the positional embeddings from *E*_pos_ : ℤ^n^ →^ℝn×d^ to include the loci spatial information. E is processed by a Multi-head Transformer, consisting of two transformer encoders, which utilizes self-attention to learn the inter-loci relations and contextualizes E. Subsequently, E is processed by *E*_str_ : ℝ^n×d^ →^ℝn×3^, consisting of a linear decoder, to predict 3D loci coordinates. These coordinates are centered and divided by the maximum loci *L*_2_ norm to produce 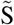. 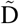 is derived from 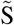 by computing pairwise Euclidean distances. A major challenge in the addressed task is the lack of ground truth 3D chromatin structures, which inhibits assessment of model predictions. As a proxy, we approximate the prediction uncertainty by producing loci-level confidence scores. Formally, we linearly project E to 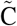, which represents the non-normalised predictions/logits for belonging to 100 confidence interval bins. 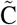 is used for confidence score optimization and model calibration as described below.

### Loss objectives

We optimize ℒ = ℒ_*d*_ + *λ*_*k*_ *ℒ*_*k*_ + *λ*_*c*_ *ℒ*_*c*_, consisting of three loss terms, to tune ChromFormer. _*d*_ is the mean squared error (MSE) between D and 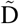 to ensure the distance preservation. The predicted structure 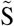 can be oriented and scaled differently with respect to S while being similar in structure. To mitigate this, we use the Kabsch algorithm [10] to align 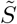 to S, resulting in 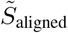. ℒ_k_ computes the MSE between S and 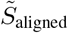. ℒ_c_ aims to ensure the correct prediction loci confidence, computed using a cross entropy between Softmax(C) and estimated confidence scores C. C is derived using a modified AlphaFold’s [9] Local Distance Difference Test [15]. First, an inverse relative error 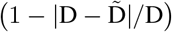 is computed to assign low confidence to large deviations between D and 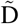. The error is row-normalized excluding self-interacting diagonal to produce the scores. C is defined as the one hot encoding of ⌊scores × 100⌋. All implementation details are given in the Appendix A.3.

### Calibration network

In the absence of real ground truth structures, it is crucial to define loci-level confidence scores. ℒ_*c*_ is weighted low during model training to give more emphasize to accurate structure prediction that induces sub-optimal confidence 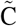. We design a calibration network to correct for this and improve confidence estimates on unseen real test data. Additionally, the network can also correct for overconfident predictions in 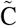. To train the calibration network, we use C and 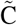 from the synthetic validation set *v*, which are then split into 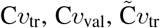, and 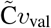 for network training and validation. We train three networks supporting three calibration techniques, *i*.*e*., temperature scaling [7], isotonic calibration [8], and beta calibration [11]. The networks transform 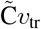 to calibrated 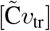 to match 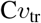. The performance of the networks is quantified by, MSE(calibrated 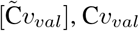 − MSE 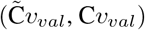. Upon training, these networks are applied on 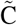, predicted by ChromFormer, for the real test data.

## 3 Results and Discussion

### Public Datasets

We exploit the following publicly available datasets: the **T****russart** **dataset** [23], a set of 100 simulated structures that contain loops, TADs and long-range interactions paired with a single Hi-C matrix (202 loci at 5 kilobase resolution), and the **T****anizawa** **dataset** [22], a real experimental single Hi-C measurement across the full fission yeast genome that contains 1258 loci at 5 kilobase resolution together with 18 ground truth pair-wise 3D Euclidean distances as measured by Fluorescence In Situ Hybridization (FISH).

### Synthetic dataset

Using the Uniform Cluster Walk algorithm, we generated 1000 and 500 structures with 202 and 1258 loci to match the Trussart and Tanizawa datasets, respectively (see Appendix Figure 4). We follow a 4:1 train-validation split for training ChromFormer. The validation set is further split into a 9:1 ratio for the calibration.

### Evaluation metrics

For the Trussart data, the predicted 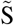 by ChromFormer is aligned with the {S_*i*_}_*i*=1:100_ ground truth structures using the Kabsch algorithm and the mean of 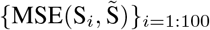 is used to quantify the model performance. For the Tanizawa data, we match the experimentally determined 18 ground truth distances to the equivalent entries in the predicted 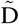, and compute the Pearson correlation between the two sets. To quantify the calibration networks, we compute MSE(C, Softmax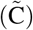) on the Trussart dataset. ReLU 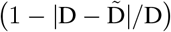 is row-normalized to get C, where D is derived from the mean of the Trussart structures.

### Qualitative and quantitative analysis

Figure 2 displays a synthetic structure S and its corresponding Hi-C matrix H. We clearly observe that S is smooth with equidistant consecutive points and that it includes chromatin loops and TAD-like domains, that are also visible in H. The Hi-C matrices in Figure 2(b) and (e) highlight the distribution gap between the synthetic and the real Trussart data, which is progressively reduced following ICE (c) and OT (d), leading to a final Hi-C matrix that is closer to the real data distribution while retaining the initial Hi-C information. Appendix Figure 5 presents more sample examples. Figure 3 presents the ground-truth Trussart structure (mean of 100 simulations) and the structure as predicted by ChromFormer using the Trussart Hi-C. We observe that the predicted structure greatly resembles the real one as it preserves all loops and TADs. Predicted structures by the competing methods are presented in Appendix Figure 7.

**Figure 2:**
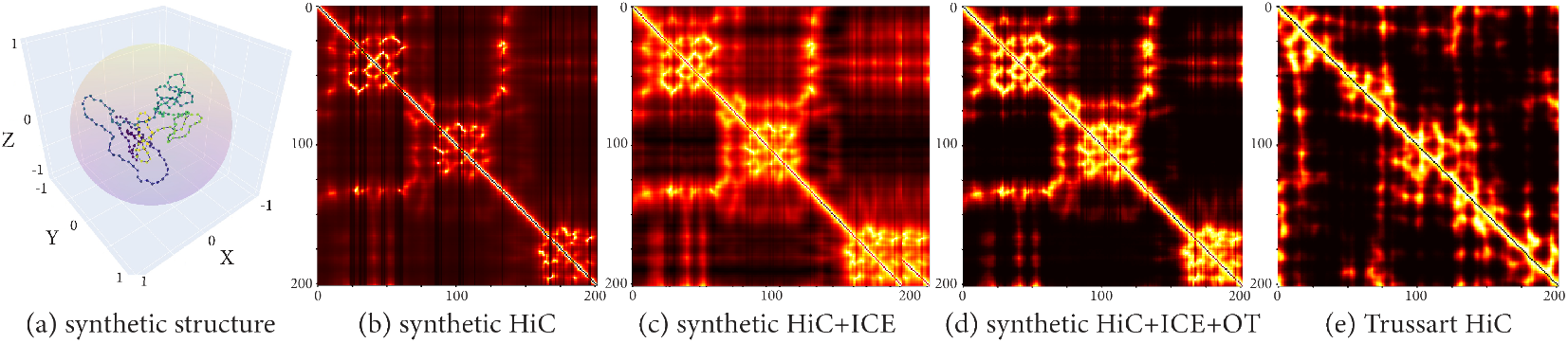
Synthetic structure (a) and Hi-C (d), and its evolution (b-d) w.r.t Trussart Hi-C (e).

**Figure 3:**
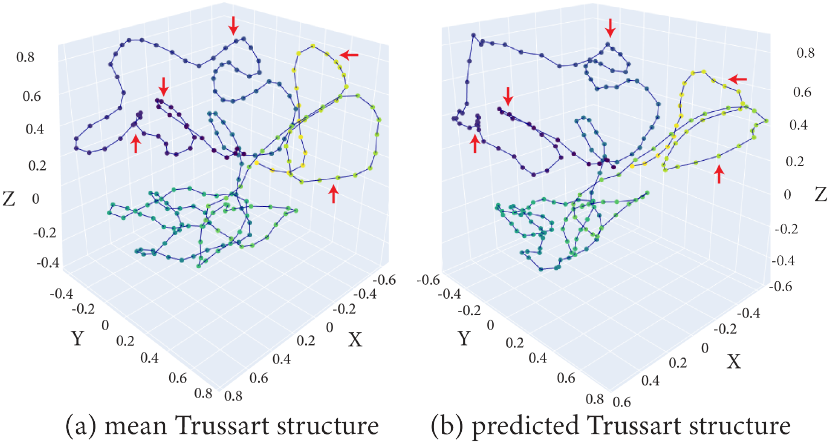
Mean ground truth vs. predicted Trussart structure. Arrows (in red) show sample regions where loops and TADs are preserved.

**Figure 4:**
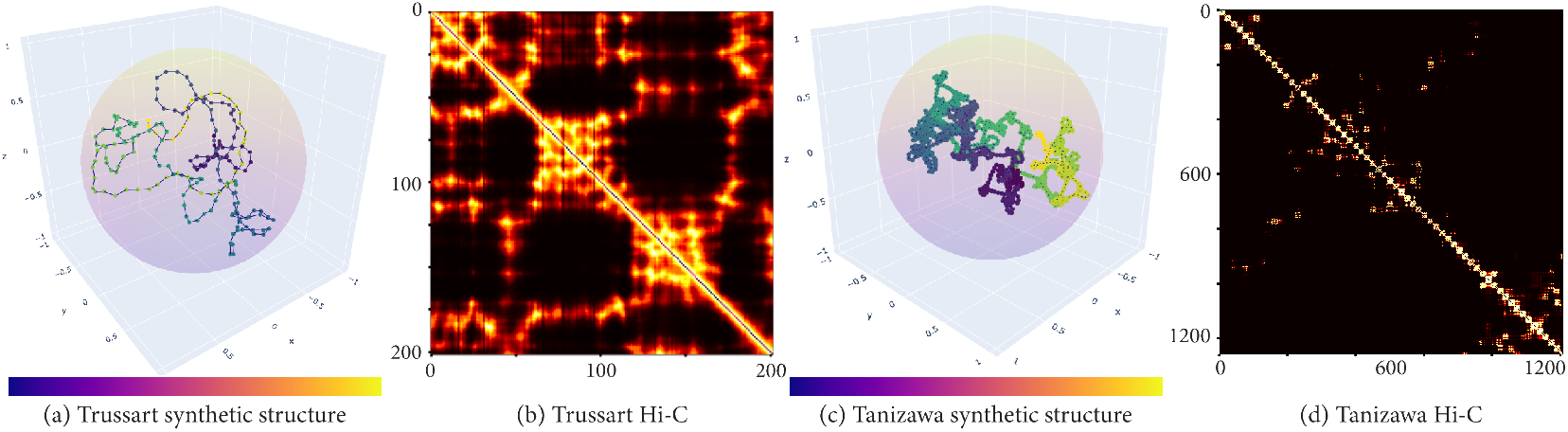
Pairs of synthetic structures and Hi-C matrices for Trussart and Tanizawa.

Quantitative results on the test datasets are presented in Table 1. ChromFormer achieves the lowest Kabsch distance on the Trussart data, outperforming competing methods by a large margin. On the Tanizawa data, ChromFormer outperforms the state-of-the-art method TECH-3D [6], with comparable performance to MiniMDS [18] and GEM [27]. Still, qualitative results (Appendix Figure 8) demonstrate that ChromFormer reconstructs smooth and loopy structures with a clear distinction between chromosomes, while miniMDS outputs a structure which is a compact set of points without loops and biological feasibility. Among the calibration networks, isotonic calibration results in the best score, followed by beta calibration. The average ground truth confidence, and the average predicted confidence before and after isotonic calibration are 88.90%, 87.40%, and 88.24%, respectively. The average uncalibrated confidence is already good, and the calibration step brings it closer to the upper bound (Appendix Table 2 and Figure 9). On the MSE metric, isotonic calibration produces a score of 8.64 compared to 10.39 before calibration.

**Table 1:** Quantitative benchmarking of our proposed method with competing algorithms on the test datasets. Best scores in **bold**.

| Methods | TRUSSART [↓]<br>(Kabsch distance) | TANIZAWA [↑]<br>(Pearson's $\rho$) |
| --- | --- | --- |
| Random | 0.2233 | -0.1044 |
| MiniMDS | 0.1439 | 0.9092 |
| GEM | 0.1437 | **0.9600** |
| REACH-3D | 0.0595 | 0.4831 |
| TECH-3D | 0.0214 | 0.7088 |
| CHROMFORMER | **0.0068** | 0.9047 |

**Table 2:** Quantitative results for the calibration networks on Trussart data. Best scores in **bold**.

| Methods | average(Softmax($\tilde{C}$)) [in %][↑] | MSE(C, $100 \times \text{Softmax}(\tilde{C})$) [↓] |
| --- | --- | --- |
| Ground truth | 88.90 | 0.00 |
| Before calibration | 87.40 | 10.39 |
| Temperature calibration [7] | 87.16 | 11.16 |
| Isotonic calibration [8] | **88.24** | **8.64** |
| Beta calibration [11] | 88.17 | 8.72 |

## 4 Conclusion and Future work

In this work we present a novel approach for reconstructing 3D genome structures that overcomes the lack of ground-truth data using a biologically-informed synthetic data generation and a transformer-based architecture. Our model can accurately reconstruct publicly available synthetic structures, outperforming state-of-the-art methods, and shows comparable performance to state-of-the-art algorithms on experimental microscopy measurements from real genomes. A limitation of our framework is data generation, a computationally intensive process that needs to be tuned to tested datasets. We are currently tuning our model hyperparameters and extending its validation to other datasets, especially single-cell Hi-C data [21].

## A. Appendix

### A.1 Uniform Cluster Walk algorithm

#### Algorithm 1: Synthetic structure generation algorithm

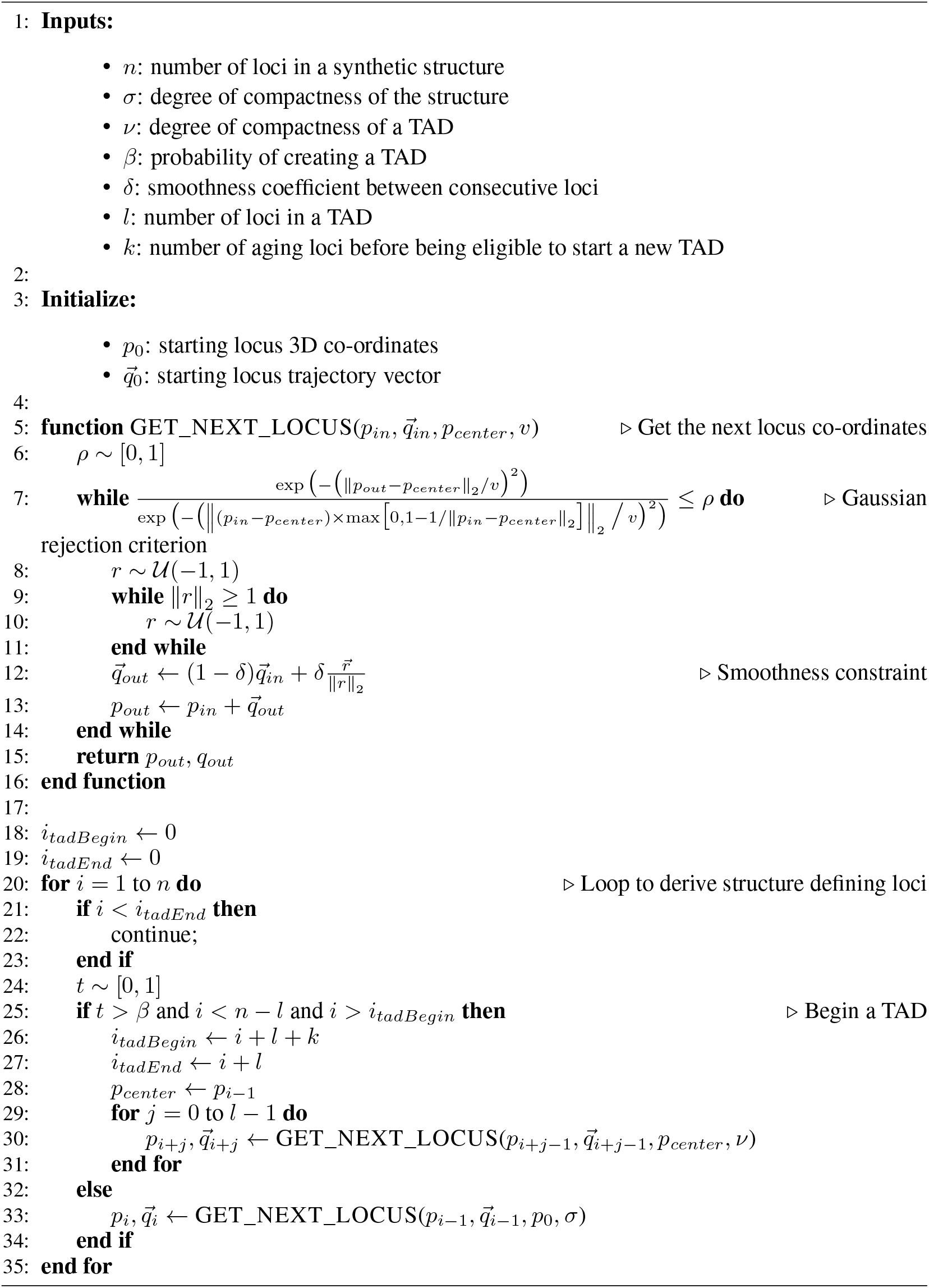

### A.2 Optimal transport

We apply OT as an unsupervised domain adaptation method to minimize the differences in values between the synthetic and the real Hi-C matrices by matching the frequency distribution of the synthetic and real values. Figure 5 shows the histograms of the synthetic Hi-C values before and after applying OT. In particular, given a set of synthetic and real Hi-C matrices, we first compute sequences of 500 sampled values from their distribution. Then, sequences are normalized to have a total mass of 1, so that they can be considered discrete probability distributions. Given the two normalized sequences *α, β*, their supports *A, B* with equal size (|*A*| = |*B*| = ℝ), and squared euclidean distance costs *c* : *A* × *B* → ℝ; *c*(*a, b*) = (*a* − *b*)^2^, the OT problem is defined as follows:

**Figure 5:**
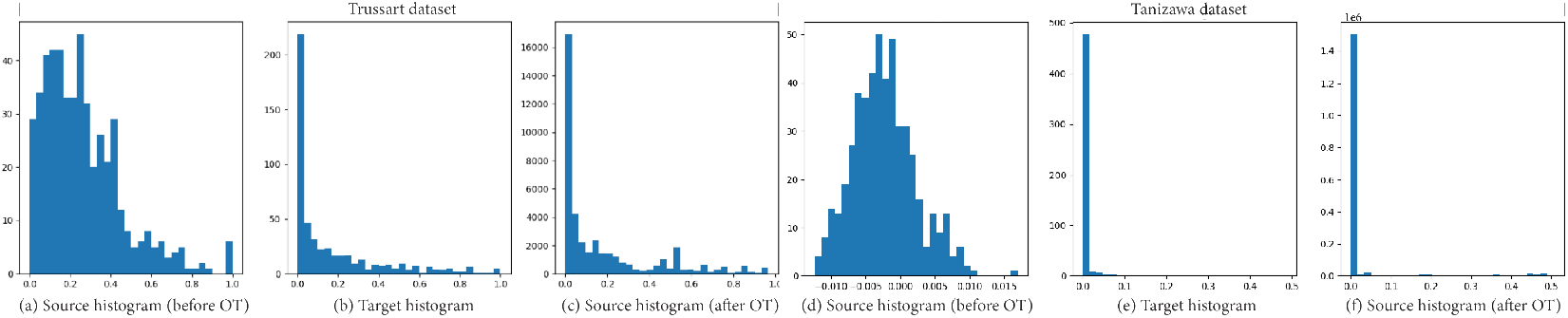
Frequency distribution of Hi-C values for Trussart and Tanizawa datasets.

**Figure 6:**
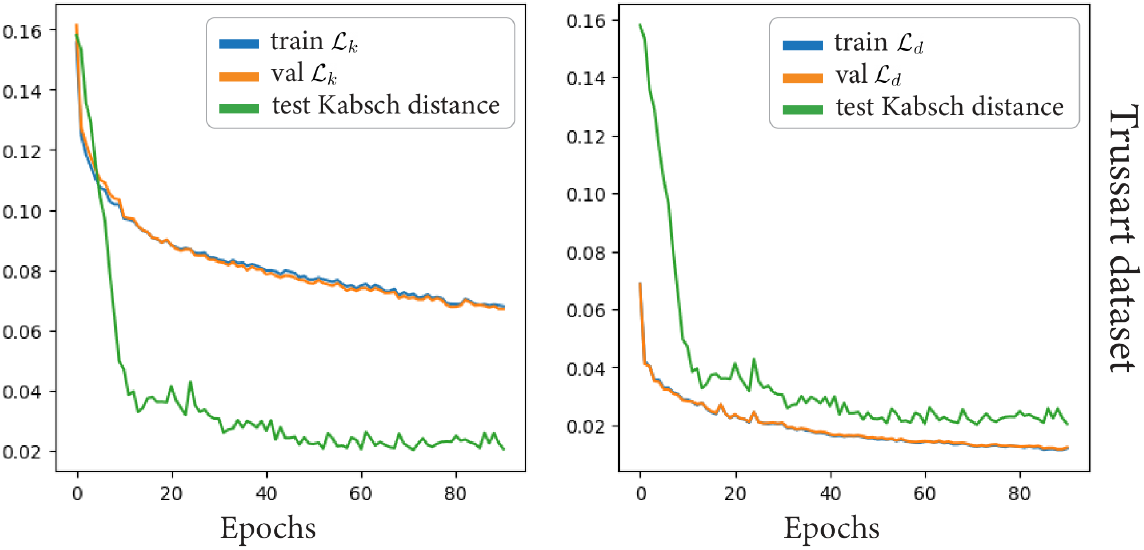
Evolution of ℒ_*k*_ and ℒ_*d*_ losses (train and validation), and evolution of Kabsch distance on test during model training on Trussart.

**Figure 7:**
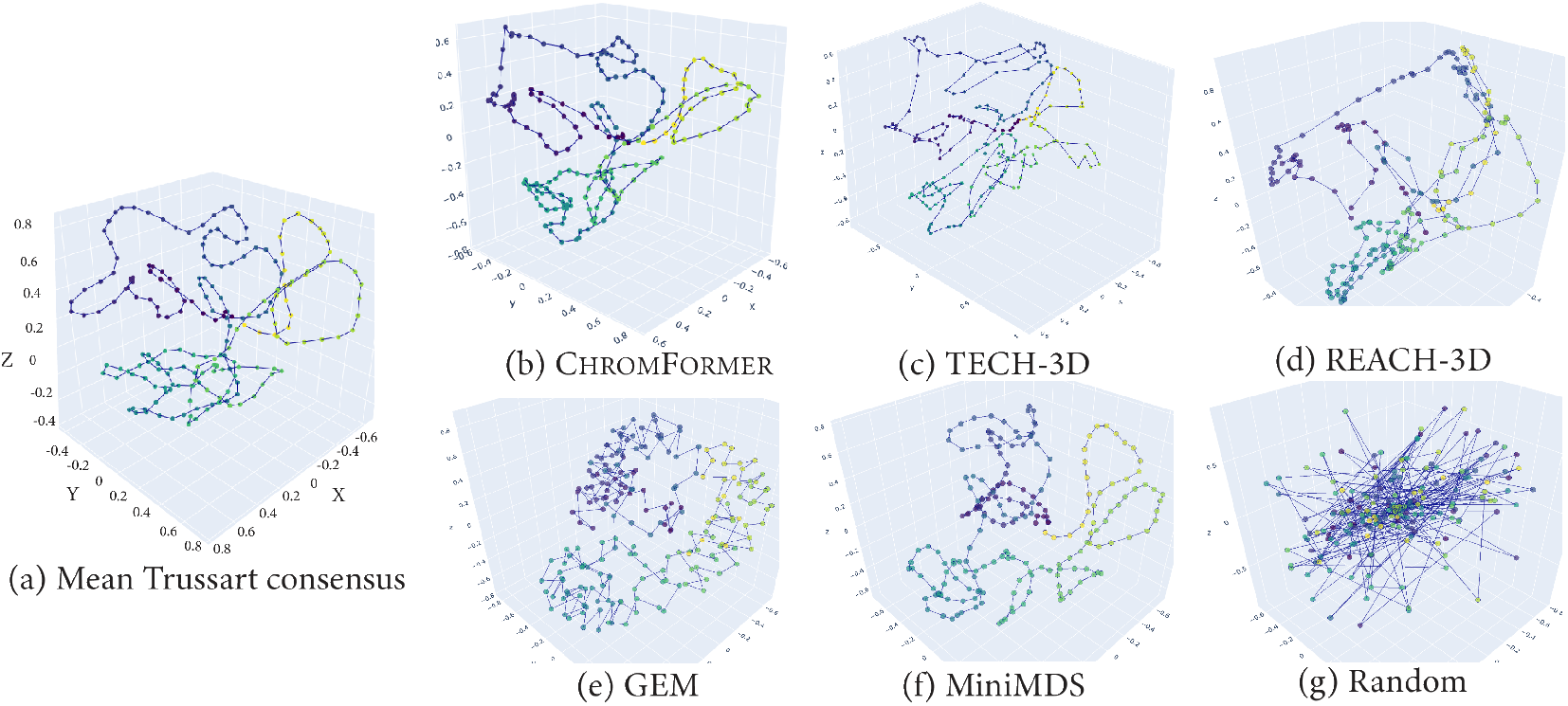
Predicted chromatin structures on Trussart by different algorithms.

**Figure 8:**
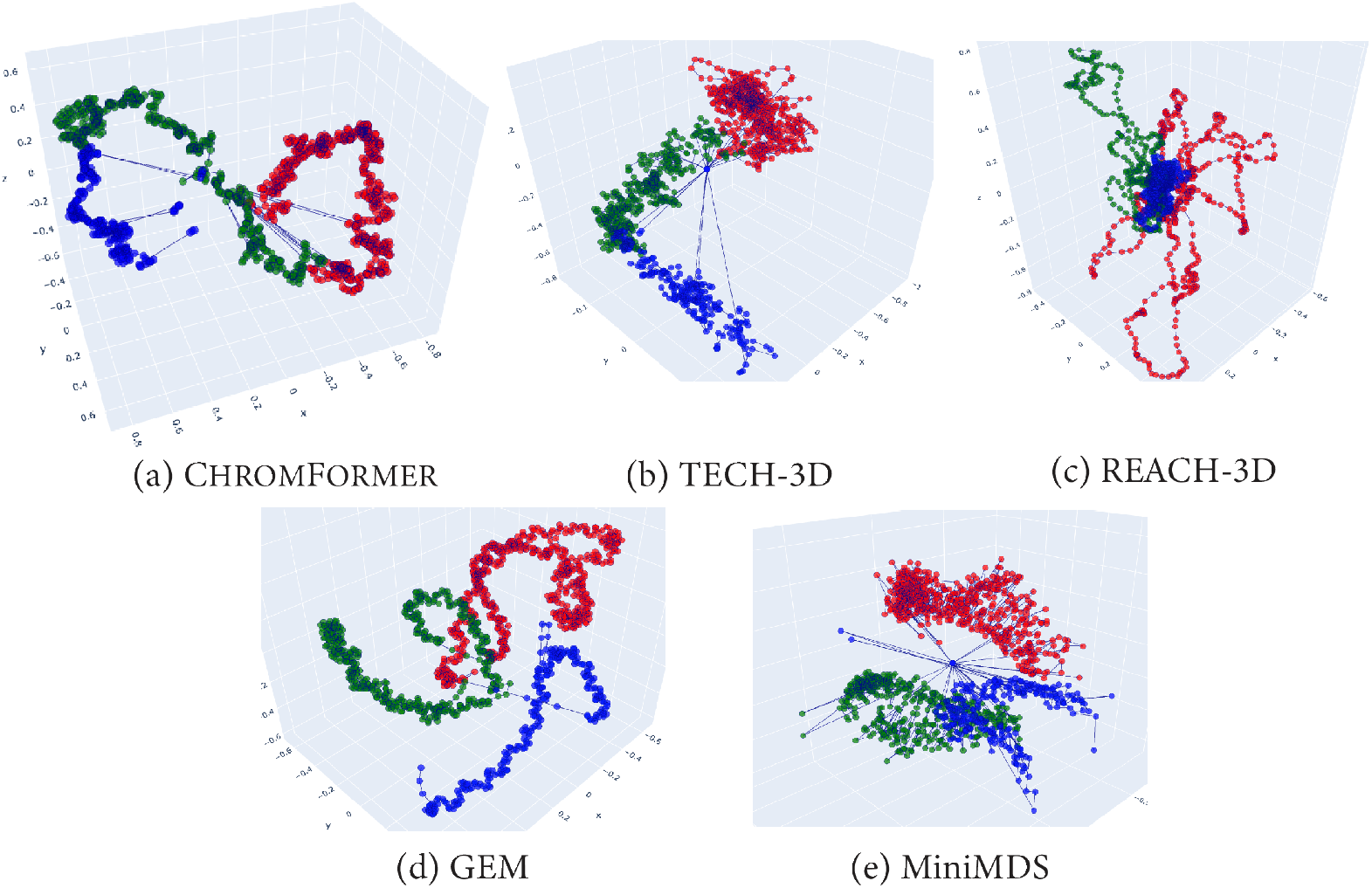
Predicted chromatin structures on Tanizawa by different algorithms.

**Figure 9:**
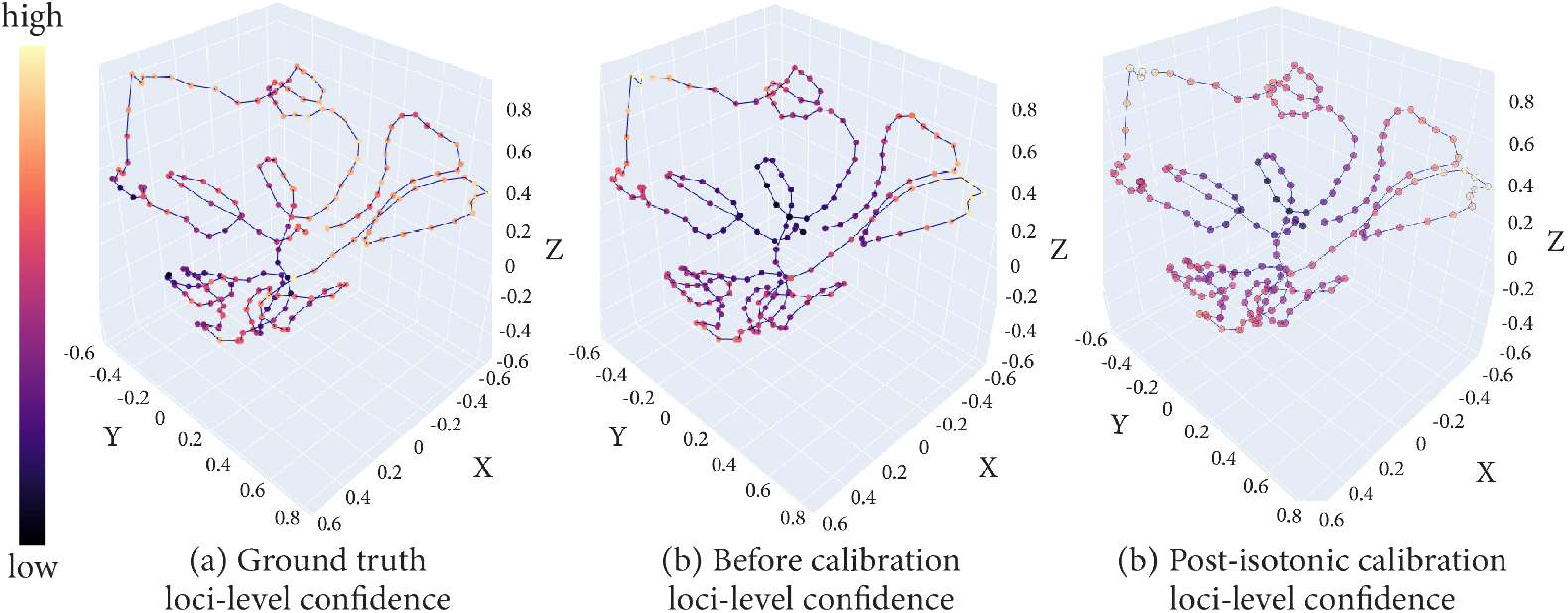
Heatmaps of loci-level confidence on Trussart dataset.

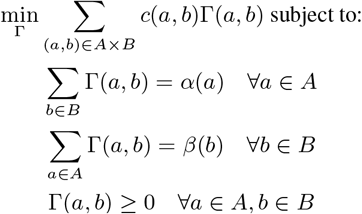

The solution of the above linear program is the optimal transport plan Γ that maps each bin *a* ∈ *A* of the source sequence *α* to one or more values *b* ∈ *B* of the target sequence *β*. The value of each Γ(*a, b*) denotes the amount of transported mass from *α*(*a*) to *β*(*b*). Finally, the source sequence is transported to the target sequence by using the optimal transport plan Γ. With the transported sequence, the values of the synthetic Hi-C matrices are transformed to match the frequency distribution of the values of the real Hi-C matrices. For the implementation, we used the OT solver from Python Optimal Transport [4] library. Notably, OT transformation is independent of the specific structure of the target Hi-C, and only relies on the frequency distribution of values in the target Hi-C matrices. Figure 2 shows the effect of the OT transformation with source Hi-C (c), target Hi-C (e), and transported source Hi-C (d).

### A.3 Implementation details

On a local CPU, it takes 30 minutes to generate the Trussart synthetic data, and 6 hours to generate the Fission Yeast synthetic data. To create the Trussart and Tanizawa synthetic structures, the selected parameters were:

| Parameters | TRUSSART | TANIZAWA |
| --- | --- | --- |
| # of loci $n$ | 202 | 1258 |
| smoothness $\delta$ | 0.45 | 0.2 |
| degree of compactness of a structure $\sigma$ | 4 | 6 |
| probability of creating a TAD | 0.1 | 0.8 |
| degree of compactness of a TAD $\sigma$ | 1.5 | 1 |
| # of loci per TAD $l$ | 30 | 15 |
| # of aging loci $k$ | 30 | 5 |
| power law $\alpha$ | 1 | 1 |

ChromFormer model and training parameters were the same for both the datasets:

- embedding dimension *d*: 100
- *E*_proj_: number of layers = 1, hidden dimensions = 100
- # transformer blocks: 2 (dimension of the feedforward networks 100 and 48, respectively)
- # heads in a transformer block: 2
- *E*_str_: number of layers = 1, hidden dimensions = 100
- *E*_conf_: number of layers = 1, hidden dimensions = 100
- *E*_cal_: number of layers = 1, hidden dimensions = 100
- Training: #epochs: 100, batch size: 10, optimized: Adamw with lr: 0.0005, weight decay: 1e-5
- Loss weights: *λ*_*k*_ = 0.1 and *λ*_*c*_ = 0.1

Training on a local CPU takes 1 and 5 hours for Trussart and Tanizawa, respectively. Training on a NVIDIA P100 GPU with POWER9 CPU takes 20 and 50 minutes for Trussart and Tanizawa datasets, respectively.

### A.4 Quantitative and qualitative results

